# Extremely Drug-Resistant *Pseudomonas aeruginosa* ST309 Harboring Tandem GES Enzymes: A Newly Emerging Threat in the United States

**DOI:** 10.1101/535302

**Authors:** Ayesha Khan, Truc T. Tran, Rafael Rios, Blake Hanson, William C. Shropshire, Zhizeng Sun, Lorena Diaz, An Q. Dinh, Audrey Wanger, Luis Ostrosky-Zeichner, Timothy Palzkill, Cesar A. Arias, William R. Miller

**Author notes:** Address correspondence to William Miller.

## Abstract

Two ST309 *Pseudomonas aeruginosa* clinical isolates resistant to carbapenems and newer β-lactam/β-lactamase inhibitor combinations were found to harbor the extended spectrum beta-lactamases GES 19 and 26 genetically clustered in tandem. The combination of ceftazidime/avibactam plus aztreonam achieved cure in one patient. Phylogenetic analysis suggests that ST309 *P. aeruginosa* carrying tandem GES are an emerging lineage in the United States.

## Introduction

The Centers for Disease Control and Prevention has identified multidrug-resistant (MDR) *Pseudomonas aeruginosa* as a serious threat and treatment of such isolates often requires the use of drugs with significant toxicities [1]. Carbapenem resistance in *P. aeruginosa* in the USA is mostly mediated by non-carbapenemase mechanisms [2]. In response, novel therapeutics such as ceftolozane/tazobactam (C/T), which is stable to the pseudomonal AmpC β-lactamase and less susceptible to porin loss and drug efflux, have entered the clinic to combat this threat [3]. Although C/T remains broadly active against most clinical isolates of carbapenem-resistant *P. aeruginosa*, resistance associated with mutations in AmpC or the expression of acquired β- lactamases has been described [4–6].

The Guiana Extended Spectrum β-lactamase (GES) enzyme was first isolated from a *Klebsiella pneumoniae* obtained from a rectal swab of an infant in Cayenne, French Guiana [7] and, since then, 32 variants have been identified. In general, these enzymes confer resistance to penicillins, including ureidopenicillins, and oxyimino-cephalosporins, but show less activity against aztreonam and imipenem [7, 8]. Nonetheless, specific substitutions can significantly alter this susceptibility profile, including G243A, which improves activity against aztreonam, and G170S, conferring increased carbapenem hydrolyzing activity [9]. These enzymes are often found in association with class 1 integrons, a gene cassette acquisition system known to harbor multiple antimicrobial resistance determinants associated with mobile genetic elements [10]. A large percentage of carbapenem-resistant *P.aeruginosa* from Mexico have been associated with GES enzymes carried by class 1 integrons, with a study of isolates from Mexico City describing ST309 as a potential high risk clone [11, 12]. Here, we report the identification of two isolates of extremely drug-resistant *P. aeruginosa* ST309 causing bloodstream infections in unrelated patients and carrying simultaneously two variants of *bla*_*GES*_ within a class 1 integron. The isolates exhibited resistance to all β-lactams including novel β-lactam/β-lactamase inhibitor combinations. Phylogenetic analyses suggested that this MDR lineage is closely related to ST309 isolates found in Mexico with the potential to disseminate.

## Methods

### Bacterial Strains and Growth Conditions

Clinical *Pseudomonas aeruginosa* isolates PA_HTX1 and 2 were purified on MacConkey agar. Single colonies were tested to ensure they retained the resistance phenotype and stocks were frozen in Brucella broth plus 15% glycerol and stored at −80° C. *Escherichia coli* TG-1 was grown on Luria-Bertani (LB) agar or LB broth supplemented with 100 µg/ml ampicillin or 25 µg/mL chloramphenicol when needed. All bacteria were grown at 37° C and with gentle agitation for liquid media.

### Genetic manipulation of *bla* genes

genes *bla*_*OXA-2*_, *bla*_*GES-19*_, *bla*_*GES-26*_, and both *bla*_*GES*_ genes in combination were cloned into the isopropyl β-D-1-thiogalactopyranoside (IPTG) inducible *E. coli* expression vector pBA169 [13] and transformed into *E. coli* TG-1. Genomic DNA isolated from PA_HTX1 was used as a template, and primers are listed in **Supplementary Table 1**. Insert DNA and plasmid pBA169 were digested with EcoRI and BamHI (New England Biolabs) and ligated overnight before transformation into chemically competent *E. coli* TG-1. Transformants were screened on LB agar containing chloramphenicol plus ampicillin. Candidates were verified by PCR, restriction digests and Sanger sequencing to confirm the presence of the *bla* inserts.

**Table 1.**
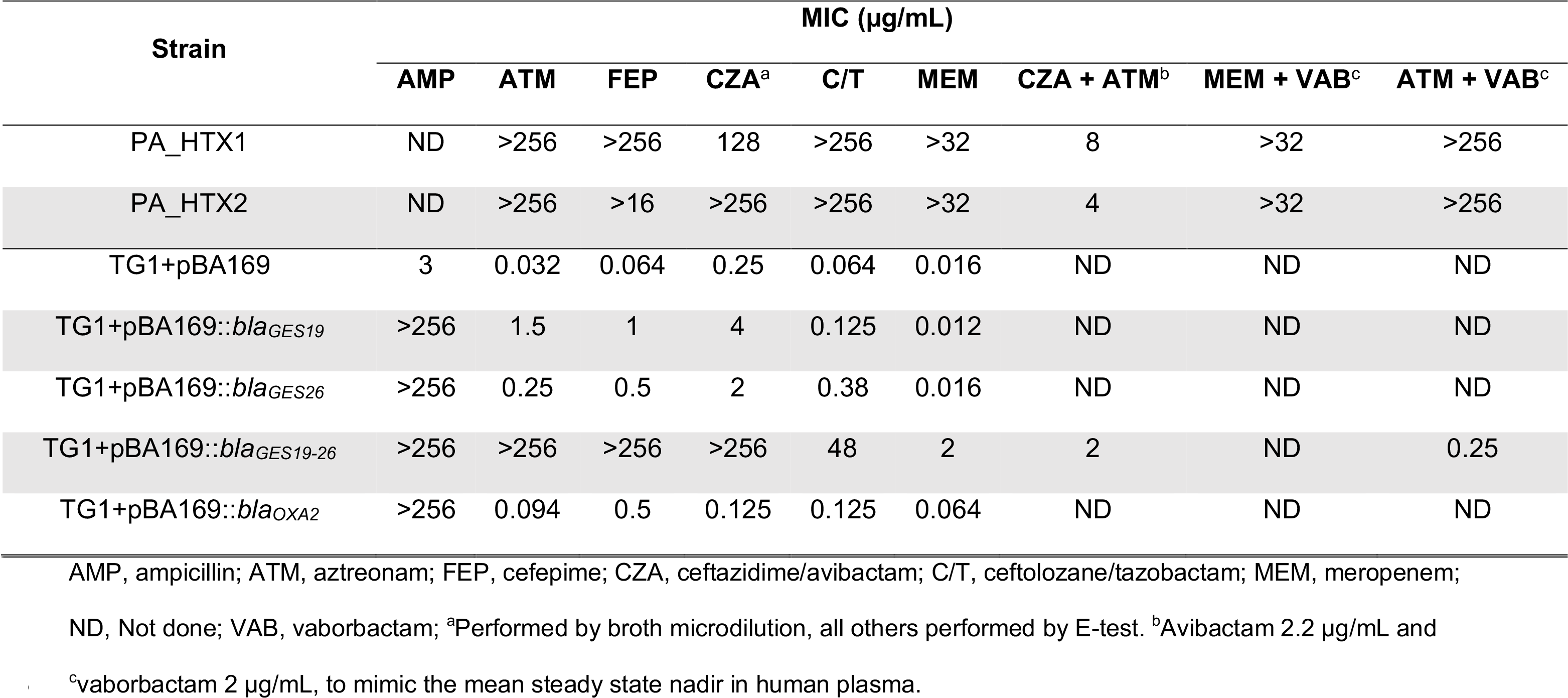
Minimum Inhibitory Concentrations for β-lactams and β-lactam/β-lactam inhibitor combinations.

| Strain | MIC (μg/mL) | | | | | | | | |
| --- | --- | --- | --- | --- | --- | --- | --- | --- | --- |
|  | AMP | ATM | FEP | CZA<sup>a</sup> | C/T | MEM | CZA + ATM<sup>b</sup> | MEM + VAB<sup>c</sup> | ATM + VAB<sup>c</sup> |
| PA_HTX1 | ND | >256 | >256 | 128 | >256 | >32 | 8 | >32 | >256 |
| PA_HTX2 | ND | >256 | >16 | >256 | >256 | >32 | 4 | >32 | >256 |
| TG1+pBA169 | 3 | 0.032 | 0.064 | 0.25 | 0.064 | 0.016 | ND | ND | ND |
| TG1+pBA169::*bla*<sub>GES19</sub> | >256 | 1.5 | 1 | 4 | 0.125 | 0.012 | ND | ND | ND |
| TG1+pBA169::*bla*<sub>GES26</sub> | >256 | 0.25 | 0.5 | 2 | 0.38 | 0.016 | ND | ND | ND |
| TG1+pBA169::*bla*<sub>GES19-26</sub> | >256 | >256 | >256 | >256 | 48 | 2 | 2 | ND | 0.25 |
| TG1+pBA169::*bla*<sub>OXA2</sub> | >256 | 0.094 | 0.5 | 0.125 | 0.125 | 0.064 | ND | ND | ND |

AMP, ampicillin; ATM, aztreonam; FEP, cefepime; CZA, ceftazidime/avibactam; C/T, ceftolozane/tazobactam; MEM, meropenem;
ND, Not done; VAB, vaborbactam; <sup>a</sup>Performed by broth microdilution, all others performed by E-test. <sup>b</sup>Avibactam 2.2 μg/mL and
<sup>c</sup>vaborbactam 2 μg/mL, to mimic the mean steady state nadir in human plasma.

### Antimicrobial Susceptibility Testing

Initial antimicrobial susceptibility testing of the clinical isolates PA_HTX1 and 2 was performed in the clinical laboratory using a Microscan Walk-Away and E-test (for colistin, C/T and ceftazidime/avibactam [CZA]). Synergy testing of the *Pseudomonas aeruginosa* isolates was performed by applying an aztreonam (ATM) or meropenem (MEM) E-test strip to Mueller-Hinton agar plates containing either CZA (Allergan), at a final concentration of 2.2 µg/mL of the avibactam component, or vaborbactam (The Medicines Co.) at 2 µg/mL. This concentration was selected to mimic the serum nadir of avibactam and vaborbactam from human PK/PD data. Antimicrobial susceptibility testing for the *Escherichia coli* TG-1 mutants was performed in triplicate by inducing strains with 1 mM IPTG for 2 hours prior to plating on Mueller-Hinton agar plates previously spread evenly with 40 µL of 100 mM IPTG. E-test strips (Biomerieux) for each antimicrobial were applied and MICs were read after 24 hours of incubation at 37° C. MICs for CZA alone were determined by broth microdilution with a standard inoculum prepared from the induced cultures into fresh MH broth without IPTG.

### Whole genome sequencing and *P. aeruginosa* phylogenetics

Genomic DNA was extracted from 2 mL of overnight growth in BHI broth using the DNeasy Blood and Tissue kit (Qiagen). Genome sequencing of the two isolates was performed on a Miseq platform using Illumina technology, and with MinION (Oxford Nanopore Technologies, United Kingdom) for long reads. Sequence data have been deposited in the NCBI database (Bioproject: PRJNA414583). Resistance detection, polymorphism analysis, and reconstruction of the phylogenetic tree were performed using a custom pipeline. A detailed description of the analysis can be found in the supplementary material. To study the phylogenetic relationships between the isolates with other *P. aeruginosa* genomes, all assembled genomes of *P. aeruginosa* available at the NCBI genome database were downloaded and the MLST was obtained using the mlst tool. Nine additional ST309 genomes were identified, and PAO1 (GCA_000006765.1), PA_D1 (GCA_001721745.1), L10 (GCA_002223805.1), M18 (GCA_000226155.1) and FRD1 (GCA_000829885.1) from the STs 539, 1971, 253, 1239 and 111 respectively, were used as references (**Supplementary Fig. 1**).

**Figure 1.**
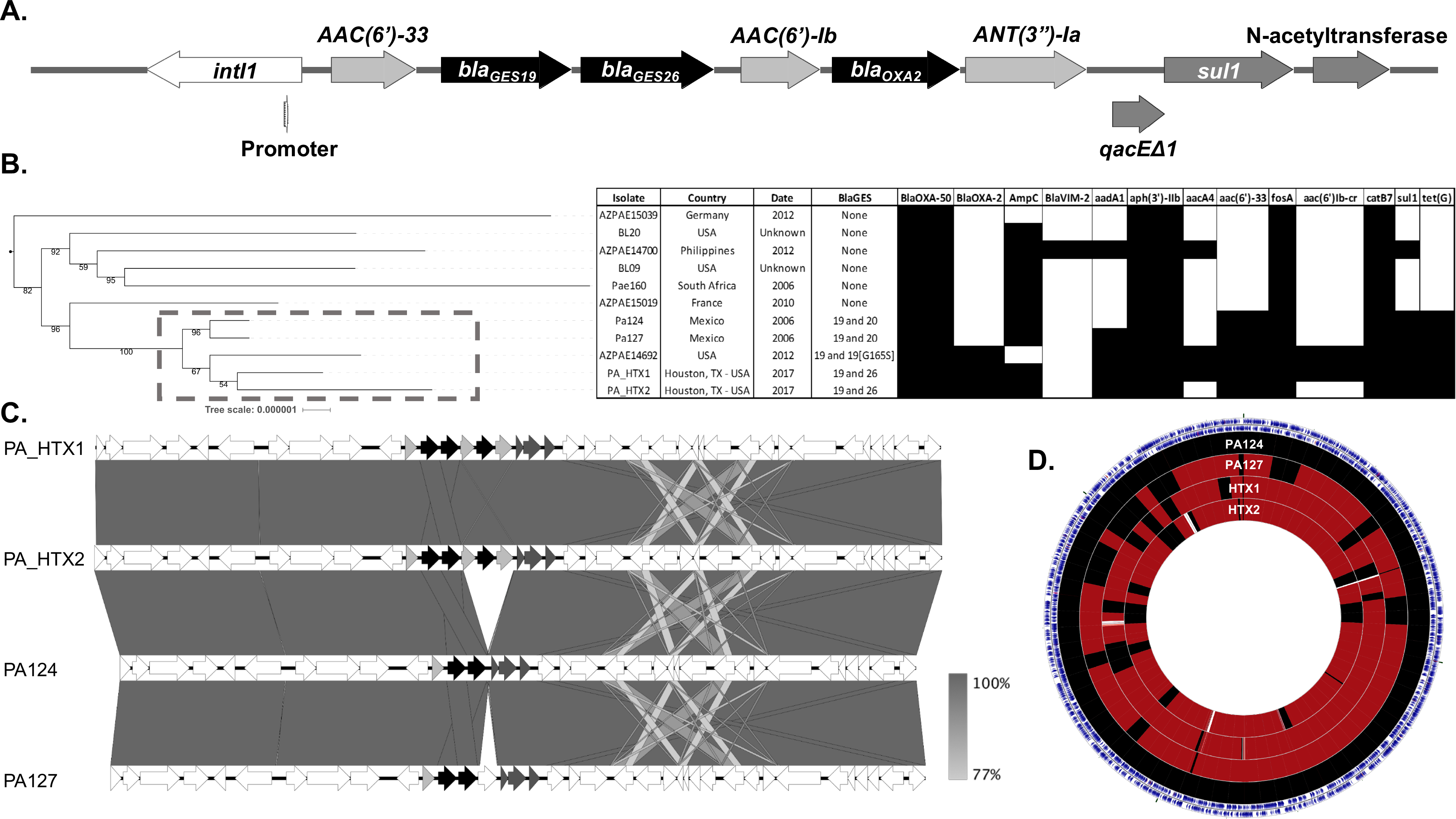
Integron structure, phylogenetics and resistome of ST309 *Pseudomonas aeruginosa* isolates available in the NCBI database. (A) Structure of the class 1 integron in PA_HTX1. Gene names are listed next to the predicted open reading frames (arrows). The integrase (white) and internal promoter, aminoglycoside modifying enzymes (AME, light gray), beta-lactamases (black), and other predicted resistance genes (dark gray) are shown. (B) A core genome based tree (RAST annotations) of ST309 using the reference genomes PAO1, PA_D1, L10, M18 and FDR1. The root of the tree was defined before the split of ST309, the clade of the references was removed and distance were set to allow resolution of the ST309 branch lengths (shown to the left, see Supplemental Figure 1 for complete tree). Isolate, origin, year, GES enzyme type and presence (black box) or absence (white box) of resistance genes is shown to the right. A gray box with dashed line indicates the GES positive isolates. (C) Comparison of the genetic context of *bla*_*GES*_ between the Houston and Mexican isolates. In addition to the acquisition of OXA-2 and AMEs, there is a downstream region of variability associated with an IS*6* transposon mobile genetic element. Grayscale gradient bar denotes nucleotide sequence identity. (D) Sequence based alignment of the genomes of the Houston and Mexican isolates using PA124 as the reference. Outer ring of blue arrows represents predicted open reading frames. Regions in black represent 100% nucleotide identity on BLAST hits, regions colored maroon represent at least 98% identity. Areas represented in white show gaps in the alignment, associated with presence or absence of phage or mobile genetic elements.

## Results and Discussion

Two patients admitted to separate hospitals in a large urban hospital network developed bacteremia due to MDR-*P. aeruginosa*. In the first case, the patient had two weeks of prior exposure to cefepime, metronidazole, and vancomycin, then meropenem for a complicated intra-abdominal infection before isolation of a MDR-*P. aeruginosa* (PA_HTX1, **Table 1 and Supplementary Table 2**) resistant to all β-lactams, including novel β-lactam/β-lactamase inhibitor combinations. Despite therapy with polymyxin B, meropenem, and amikacin, the patient’s clinical condition worsened. On the suspicion that the isolate may harbor a metallo-β- lactamase, synergy testing with the combination of ceftazidime/avibactam plus aztreonam was performed, with an observed reduction in the aztreonam MIC from >256 µg/mL alone to 8 µg/mL in the presence of avibactam. The patient was started on ceftazidime/avibactam 1.5 grams given after hemodialysis with aztreonam, 2 grams IV daily and was ultimately cured with complete resolution of abdominal abscesses after 105 days of therapy. In the second case, the patient had bronchoalveolar lavage cultures positive for *Escherichia coli*, and received five days of meropenem, before developing MDR-*P. aeruginosa* pneumonia with a similar antimicrobial susceptibility profile to PA_HTX1 (PA_HTX2, **Table 1 and Supplementary Table 2**). The patient received colistin, meropenem and three doses of ceftolozane/tazobactam, but developed bacteremia despite antibiotics and ultimately died due to the infection. An epidemiologic investigation within the hospital system did not reveal a link between the two patients, and there was no shared intensive care unit staff or equipment between the two hospitals during the period of patient admissions.

We performed whole genome sequencing on PA_HTX1 and 2. Both isolates were identified as sequence type 309 and carried a chromosomal class 1 integron with multiple resistance determinants, including *bla*_*GES-19*_ and *bla*_*GES-26*_ (**Fig. 1A, Supplementary Table 3**). To determine if this tandem carriage of GES β-lactamases was present in other ST309 *P. aeruginosa*, we performed a search of the assembled *P. aeruginosa* genomes available in the NCBI database. Eleven ST309 *P. aeruginosa* (including the two patient isolates in this study) were identified. Interestingly, a cluster of five ST309 isolates of clinical origin harboring tandem *bla*_*GES*_ genes from the United States and Mexico, (isolated from 2006 to 2017), appear to form a distinct group (**Fig. 1B**). Indeed, GES enzymes seem to be an important contributor to carbapenem resistance in *P. aeruginosa* from Mexico with a prevalence of 30.6% [11]. An analysis of the integron and surrounding genomic context revealed the Houston isolates possessed an OXA-2 β-lactamase and aminoglycoside modifying enzymes not present in the Mexican isolates (**Fig. 1C**). In addition, the strains possess a high degree of sequence identity, suggesting a close relationship between the isolates, although several gaps in the genomic alignment of the Houston strains are likely driven by phage and mobile genetic elements (**Fig 1D**).

To investigate the genetic bases of the MDR-phenotype, we performed a single nucleotide polymorphism analysis of genes associated with antimicrobial resistance using *P. aeruginosa* PAO1 as the reference strain (**Supplementary Table 4**). The *ampC* gene for both PA_HTX strains codes for the PDC-19a variant, and the polymorphisms identified in *ampR, ampD*, and *ampG* have been previously reported in both wild type and resistant strains. The sequences of the *ampD* homologues *ampDh2* and *ampDh3*, as well as *dacB*, which encodes a low molecular weight penicillin-binding protein associated with AmpC expression, were identical to PAO1. The *oprD* gene was disrupted by insertion of ISPa*1328*, an IS*256* family element that truncated the first 126 base pairs of *oprD* including the start codon. This is predicted to result in loss of *oprD* expression and likely contributes to the carbapenem resistance phenotype seen in these isolates. In addition, both HTX isolates possessed a mutation predicted to result in a deletion of 6 amino acids (residues 189-194) near the C-terminal end of MexZ, the repressor of the MexXY efflux pump. This region lies at the dimerization interface of MexZ, and in this context we hypothesize that loss of these residues may lead to upregulation of the MexXY efflux pump [14]. Mutations in *gyrA* and *parC* seen in these isolates have been previously linked to decreased susceptibility to fluoroquinolone antibiotics [2].

The PA_HTX isolates were resistant to all available β-lactams, including the novel combinations of C/T, CZA and meropenem/vaborbactam (**Table 1**). In the presence of CZA (2.2 µg/mL avibactam component), the MIC of ATM decreased from >256 µg/mL in PA_HTX1 and PA_HTX2 to 8 and 4 µg/mL, respectively. No change in MIC was found with either meropenem or aztreonam in combination with vaborbactam. In the case of meropenem, disruption of OprD and the resulting decrease in permeability to meropenem would not be overcome with addition of a β-lactamase inhibitor. The lack of efficacy when paired with ATM is likely multifactorial, as vaborbactam is a less potent inhibitor of class C β-lactamases, does not inhibit class D enzymes, and may possibly be impacted by porin mutations (vaborbactam uses OmpK36 and OmpK35 to cross the membrane in *Klebsiella pneumoniae*), although data regarding vaborbactam entry in *Pseudomonas* are lacking [15]. In addition, some variants of the GES enzymes are relatively more resistant to inhibition by β-lactamase inhibitors, although avibactam appears to retain activity [16].

To evaluate the spectrum of the integron encoded β-lactamases, *bla*_*GES-19*_, *bla*_*GES-26*_, and *bla*_*OXA-2*_ from PA_HTX1 were cloned individually, and *bla*_*GES-19*_ and *bla*_*GES-26*_ in tandem, into *E. coli* TG-1 using an inducible vector (pBA169) that responds to isopropyl β-D-1- thiogalactopyranoside (IPTG) [13] (**Table 1**). The presence of *bla*_*GES-19*_, *bla*_*GES-26*_, and *bla*_*OXA-2*_ alone led to modest increases in MIC for ATM, cefepime (FEP), CZA (GES only) and C/T, but not MEM, as compared to TG-1 carrying empty pBA169. In contrast, the presence of both *bla*_*GES-19*_ and *bla*_*GES-26*_ resulted in a marked increase of MICs of ATM, FEP, CZA, C/T and MEM. This effect was reversed by addition of the β-lactamase inhibitors avibactam and vaborbactam. While the IPTG induced levels of enzyme are not physiologic, the results suggest that the combination of enzymes acts synergistically. Cell lysates of induced TG-1 with each of the GES constructs was tested for hydrolysis of nitrocefin and ceftazidime (**Supplemental Figure 2**). Nitrocefin hydrolysis by the cell lysate of GES-19, GES-26 and GES-19/GES-26 was 0.34 ± 0.03, 0.19 ± 0.006 and 1.34 ± 0.02 μM per sec per μl of cell lysate, respectively. Interestingly, cell lysate of the combined GES19/GES26 is 4-fold more efficient than that of GES19 alone and 7-fold more than GES26 alone. Minimal hydrolysis of ceftazidime was seen in the GES-26 cell lysate, while hydrolysis by GES-19 was 0.029 ± 0.0015 μM per sec per μl. With the GES- 19/GES-26 lysate, ceftazidime hydrolysis was 0.048 ± 0.0084 μM per sec per ul, a 1.7 fold increase. Although we are not able to quantify absolute differences in expression of GES enzymes in each lysate, which could account for differences in nitrocefin and ceftazidime hydrolysis activity, these data suggest expression of both β-lactamases provides synergistic activity against β-lactam antibiotics. Thus, the tandem acquisition of GES enzymes in combination with efflux and decreased permeability in *P. aeruginosa* has the potential to further compromise β-lactam activity.

In summary, we report the occurrence of serious infections caused by ST309 *P. aeruginosa* carrying *bla*_*GES-19*_ and *bla*_*GES-26*_ in Houston, TX. Tandem expression of the two GES enzymes under IPTG induction in *E. coli* resulted in resistance to all β-lactams including novel combinations. Ceftazidime/avibactam plus aztreonam was active *in vitro* against these two isolates and was successfully used in one patient failing polymyxin based therapy. The prevalence of GES enzymes reported in Mexico, and the close relation of the strains identified here to *P. aeruginosa* Mexican isolates suggests that ST309 may be a newly emerging high-risk lineage in the United States.

## Acknowledgements

We thank Christophe Herman for providing the pBA169-cm plasmid. Part of this work was presented at ASM Microbe 2018, Atlanta, GA. *Potential conflicts of interest*. C.A.A. has received grants from Merck, MeMed Diagnostics and Entasis Pharmaceuticals. L.O. has received grants and/or speaking and consulting honoraria from Merck, Astellas, Pfizer, Gilead, The Medicines Company, Cidara, Scynexis, Aradigm, and Bayer. W.R.M. has received grants and/or honoraria from Merck, Achaogen, and Shionogi. A.K., T.T.T., R.R., B.H., L.D., A.Q.D., W.C.S., Z.S., T.P., and A.W. report no conflict.

## Correspondence

William R. Miller, MD

6431 Fannin St. MSB 2.112, Houston, TX 77030

713-500-5495 fax

