## Supplementary Material for "Extremely Drug-Resistant *Pseudomonas aeruginosa* ST309 Harboring Tandem GES Enzymes: A Newly Emerging Threat in the United States"

### Supplemental Material

#### Methods

**Whole genome sequencing and characterization.** Genomic DNA was extracted from 2 mL of overnight growth in BHI broth using the DNeasy Blood and Tissue kit (Qiagen). Genome sequencing of the two isolates was performed on a Miseq platform using Illumina technology. Genome assembly was done using SPades v3.10.1 [1]. Identification of resistance elements was done using BlastN [2] searches against the ResFinder [3] database (version 14- Sept- 2017), selecting hits with an identity percentage higher than 90% and a coverage higher than 80%. MLST typing was done using the mlst tool (https://github.com/tseemann/mlst). The virulome characterization was done using BlastP [2] searches against the proteins annotated as virulence factors in the *Pseudomonas aeruginosa* Genome DB [4], selecting hits with an identity percentage higher than 95% and a coverage higher than 80% as positive. To close the genome and resolve the integron structure, PA_HTX1 was also sequenced on a MinION (Oxford Nanopore Technologies, United Kingdom). DNA prep without size selection was performed with the ligation sequencing kit 1D (SQK-LSK108) and run on a R9.4 flow cell for 48 hours. Raw data was base called and converted to fastq with Albacore 2.1.3 (Oxford Nanopore Technologies, United Kingdom). The genome was assembled with CANU 1.6 and individual reads covering the integron were aligned with LAST version 752 [5,6].

***Pseudomonas aeruginosa* ST309 phylogenetics.** To study the phylogenetic relationships between the isolates with other *P. aeruginosa* genomes, all assembled genomes of *P. aeruginosa* available at the NCBI genome database [7] up to 12- Sept-2017 were downloaded and the MLST was obtained using the mlst tool (https://github.com/tseemann/mlst). Nine genomes with ST309 were selected and also the references PAO1 (GCA_000006765.1), PA_D1 (GCA_001721745.1), L10 (GCA_002223805.1), M18 (GCA_000226155.1) and FRD1 (GCA_000829885.1) from the STs 539, 1971, 253, 1239 and 111 respectively (**Supplementary Fig. 1**). Genome annotation of the 16 genomes was carried out with RAST [8], a core genome was determined with Roary [9], multiple sequence alignment of the orthogroups belonging to the core genome was done with Muscle [10] and later concatenated to be used as the matrix for phylogenetic reconstruction with RAxML [11]. The best tree out of 20 runs was selected with a General Time Reversible evolution model and a Gamma model of rate heterogeneity with 100 bootstrap resampling. The tree was plotted with iTol [12] after rooting at the split between ST309 and the other references, modification of the distance between the root and the branches of ST309 was changed to a value near zero to allow the visualization of the branching of the ST309. Virulence genes were assessed using annotations from the *Pseudomonas aeruginosa* genomes database [4]. For all sequence types (STs) for which at least 5 genomes were available (n=75), the mean number of annotated virulence genes was calculated. The mean of each ST was then compared using 2-way ANOVA and Kruskal H-test, with equal mean of virulence elements between ST309 and each other ST as null hypothesis, defining statistical significance with a p-value lower than 0.05, the test was done with an in-house script using the Python module Numpy [13].

**Expression of GES proteins and preparation of cell lysate.** Overnight cultures of *Escherichia coli* cells transformed with GES19, GES26 or GES19/GES26 in the IPTG inducible pBA169 expression plasmid were diluted 1:100 in 50 ml LB medium containing 25 µg/ml chloramphenicol (to maintain the plasmid). The diluted cultures were incubated with shaking at 37ºC to OD600 of 0.8. IPTG was added to a final concentration of 0.5 mM and cells were incubated with shaking at 20ºC for another 20 hours to induce the expression of GES proteins. Cells were then pelleted by centrifugation at 6000 rpm for 20 min, and the cell pellet was resuspended in 2 ml B-PER (Thermo Fisher) containing 100 µg/ml lysozyme and 20 µg/ml DnaseI and incubated at room temperature for 15 min. Samples were then centrifuged again at 13000 rpm for 5 min to collect cell lysate in the supernatant.

**Enzyme assay.** In order to ensure the expression of GES enzymes, 50 µM nitrocefin diluted in 50 mM HEPES (pH7.4) was incubated with 1 µl cell lysate and absorbance at 482 nm (A482) was monitored on Beckman DU800 spectrometer using 1 cm cuvette. To determine hydrolysis of ceftazidime by the GES enzymes, 50 µM ceftazidime diluted in 50 mM HEPES (pH7.4) was incubated with 1 µl cell lysate and absorbance at 260 nm (A260) was monitored on Beckman DU800 spectrometer using a 1 cm cuvette.

**Supplementary Table 1 – Primers used in this study.**

| | |
|---|---|
| OXA-2_F | 5'- ccGGATCCggcattaaggaaaagttaatggc-3' |
| OXA-2_R | 5'- cgAAGCTTgttttatcgcgcagcgtc-3' |
| GES-19_F | 5'- ccGAATTCcttgtggaagattgcg-3' |
| GES-19_R | 5'- gcGGATCCctatttgtccgtgctcag-3' |
| GES-26_F | 5'-ccGAATTCccatctcaagggatcac-3' |
| GES-26_R | 5'- gcGGATCCctatttgtccgtgctcag -3' |
| GES-19_26_F | 5'- ccGAATTCcttgtggaagattgcg -3' |
| GES-19_26_R | 5'- gcGGATCCcgaattgttagacggg -3' |

**Supplementary Table 2 – Antimicrobial susceptibilities for the clinical *Pseudomonas* isolates.**

| Strain | MIC (µg/mL) | | | | | | | | | | | |
|---|---|---|---|---|---|---|---|---|---|---|---|---|
| | AMK | ATM | FEP | CAZ | CZA | C/T | CIP | CST | GEN | MEM | TZP | TOB |
| PA_HTX1 | 16 | >256 | >256 | >16 | 128 | >256 | >2 | 2 | 4 | >32 | >64 | >8 |
| PA_HTX2 | >32 | >256 | >16 | >16 | >256 | >256 | >2 | 1 | >8 | >32 | >64 | >8 |

AMK, amikacin; ATM, aztreonam; FEP, cefepime; CAZ, ceftazidime; CZA, ceftazidime/avibactam; C/T, ceftolozane/tazobactam; CIP, ciprofloxacin; CST, colistin; GEN, gentamicin; MEM, meropenem; TZP, piperacillin/tazobactam; TOB, tobramycin

#### Supplementary Table 3 – Resistance Determinants Identified by Whole Genome Sequencing.

##### PA_HTX1

| Class | Resistance gene(s) |
|---|---|
| Aminoglycoside | ***aadA1***, *aph(3')-IIb*, ***aacA4***, ***aac(6')-33*** |
| Beta-lactamase | ***blaGES-19, blaGES-26*, *blaOXA-2***, *blaOXA-50*, *blaPAO*(*ampC*) |
| Fosfomycin | *fosA* |
| Phenicol | *catB7* |
| Sulphonamide | ***sul1*** |
| Tetracycline | *tet(G)* |

##### PA_HTX2

| Class | Resistance gene(s) |
|---|---|
| Aminoglycoside | ***aadA1***, *aph(3')-IIb*, ***aac3-Ib, aac(6')-33*** |
| Beta-lactamase | ***blaGES-19, blaGES-26*, *blaOXA-2***, *blaOXA-50*, *blaPAO*(*ampC*) |
| Fosfomycin | *fosA* |
| Phenicol | *catB7, florR* |
| Sulphonamide | ***sul1*** |
| Tetracycline | *tet(G)* |

Genes listed in **bold** present in class 1 integron.

**Supplementary Table 4 – Non-synonymous Amino Acid changes of genes associated with antibiotic resistance in PA_HTX1 and PA_HTX2**

| PAO1 locus tag | Gene | AA in PAO1 | AA position | AA in PA_HTX1 | AA in PA_HTX2 | Notes | Reference |
|---|---|---|---|---|---|---|---|
| PA4110 | *ampC* | G | 1 | D | D | PDC-19a | [14] |
| | | T | 79 | A | A | | |
| | | V | 179 | L | L | AA numbering beginning after removal of the | |
| | | V | 330 | I | I | 26 AA leader peptide | |
| | | G | 365 | A | A | | |
| PA4109 | *ampR* | G | 283 | E | E | Associated with ceftazidime resistant *P. aeruginosa*, however, these SNPs have also been found | [15,16] |
| | | M | 288 | R | R | in WT strains | |
| PA4522 | *ampD* | Q | 44 | H | H | Q44H unknown significance | [17] |
| | | G | 148 | A | A | G148A common polymorphism present in both resistant and susceptible *P. aeruginosa* | |
| PA5485 | *ampDh2* | | - | | | WT sequence | |
| PA0807 | *ampDh3* | | - | | | WT sequence | |
| PA4393 | *ampG* | A | 583 | T | T | Present in 22 strains from Cabot survey | [17] |
| PA3047 | *dacB* | | - | | | WT sequence | |
| PA0424 | *mexR* | V | 126 | E | E | SNP present in sensitive strains | |
| PA3721 | *nalC* | G | 71 | E | E | Both SNPs present in sensitive strains | |
| | | S | 209 | R | R | | |
| PA3574 | *nalD* | | - | | | WT sequence | |
| PA2020 | *mexZ* | L | 138 | R | R | | |
| | | | 189-194 | Deletion of 6 AA | Deletion of 6 AA | 6 AA deletion at C-terminal end, predicted to eliminate dimerization interface based on crystal structure | [18] |
| PA5471 | *armZ* | C | 40 | R | R | Unknown significance | |
| | | L | 88 | P | P | | |
| | | S | 112 | N | N | | |
| | | I | 237 | V | V | | |
| | | V | 243 | A | A | | |

**Supplementary Table 4, cont. – Non-synonymous Amino Acid changes of genes associated with antibiotic resistance in PA_HTX1 and PA_HTX2**

| PAO1 locus tag | Gene | AA in PAO1 | AA position | AA in PA_HTX1 | AA in PA_HTX2 | Notes | Reference |
|---|---|---|---|---|---|---|---|
| PA4600 | *nfxB* | | - | | | WT sequence | |
| PA3168 | *gyrA* | T | 83 | I | I | Fluoroquinolone resistance associated mutation | [17] |
| | | | 912-913 | Deletion of 2 AA | Deletion of 2 AA | | |
| PA4964 | *parC* | S | 87 | L | L | Fluoroquinolone resistance associated mutation | [17] |
| | | S | 197 | L | L | | |

AA, Amino Acid; SNP, single nucleotide polymorphism; WT, wild type.

**Supplementary Figure 1. Phylogenetic Tree of ST309 *Pseudomonas aeruginosa* from the NCBI database.**

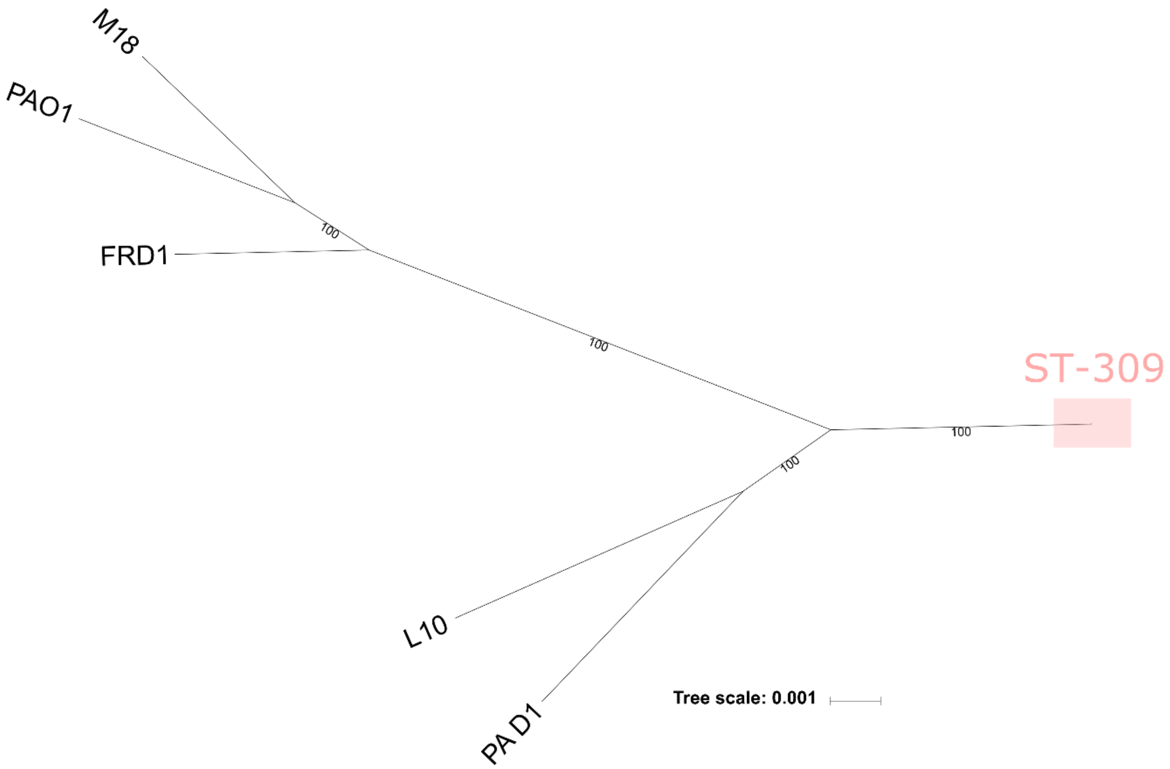


The 2637 available genomes and assemblies for *P. aeruginosa* available in the NCBI database were screened for ST309 isolates. A core genome based tree (RAST annotations) was generated with 100 bootstraps using the reference genomes PAO1, PA_D1, L10, M18 and FDR1. ST309 isolates were highly clustered (red shaded box) in relation to the outgroups shown.

**Supplementary Figure 2. Rates of hydrolysis from *E. coli* cell lysates.**

**A. Nitrocefin hydrolysis**

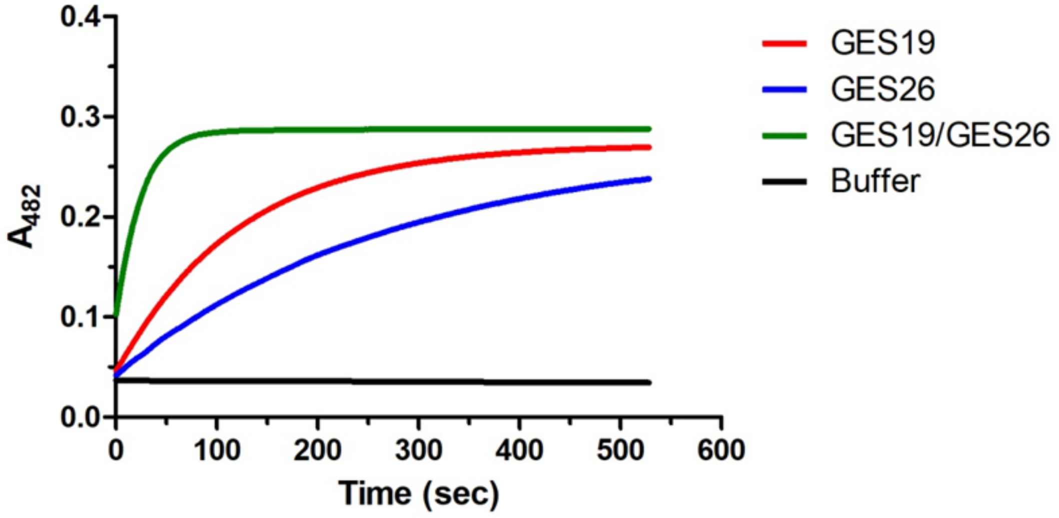


**B. Ceftazidime hydrolysis**

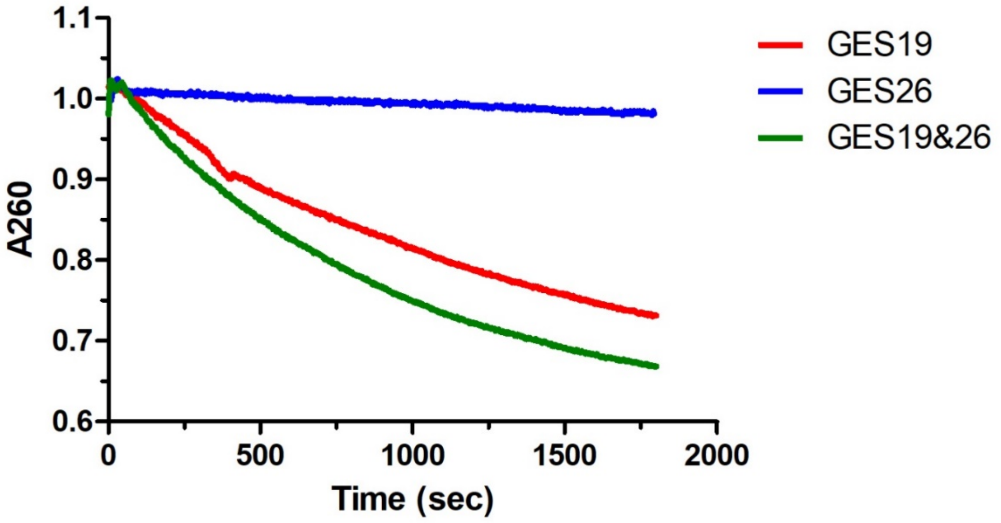


*E. coli* lysates containing GES19, GES26, and GES19/GES26 were prepared from cultures grown in the presence of 0.5 mM IPTG for 20 hours to induce protein expression. **A)** Hydrolysis of 50μM nitrocefin in 50 mM HEPES pH7.4 monitored by absorbance at 482 nm. Cell lysate with both enzymes was more efficient than that of GES19 or GES26 individually by 4-fold and 7-fold, respectively. **B)** Hydrolysis of 50μM ceftazidime in 50 mM HEPES pH7.4 monitored by absorbance at 260 nm. Hydrolysis of ceftazidime in the presence of cell lysate containing both enzymes was 1.7-fold higher than GES19 alone.
